# Baseline survey of the fish and invertebrate fauna of Hill Inlet, the northernmost estuary in south-western Australia

**DOI:** 10.1101/2020.06.21.163402

**Authors:** James R. Tweedley, Ayme Sama, Brian Poh, Neil R. Lonergan

## Abstract

Microtidal estuaries in Mediterranean climates are particularly vulnerable to the effects of anthropogenic degradation. This study provides the first data on the fish and benthic macroinvertebrate fauna of Hill Inlet, the northernmost estuary in south-western Australia. Sampling was conducted in June 2019 (Austral winter), when water levels were very high due to recent heavy rainfall and the bar at the mouth of the estuary was intact. Surface salinities were oligohaline and declined along the longitudinal axis, ranging from 12 to 3. A marked halocline was present at most sites, resulting in pronounced hypoxia. High water levels precluded the use of a seine net at some sites to sample the nearshore fish fauna, however, two species were recorded (*Pseudogobius olorum* and *Acanthopagrus butcheri*), both of which complete their life-cycle within the estuary. Deeper, offshore waters, sampled using gill nets, yielded only four species (*Mugil cephalus, A. butcheri, Adrichetta forsteri* and *Pomatomus saltatrix*), due to the bar at the mouth of the estuary being closed prior to sampling thus limiting recruitment from marine species. Ten benthic macroinvertebrates species were collected, representing mainly polychaetes, molluscs and crustaceans. The low number of species was likely caused by the hypoxia present throughout most of the bottom waters. Although these data represent a benchmark against which future changes can be detected, it is recommended that additional sampling is conducted when water levels are lower and the bar has been open to provide a more holistic assessment of the fauna of Hill Inlet.

## Introduction

Estuaries are crucial habitats providing a range of ecosystem services, including functioning as nursery areas for a range of fish and crustacean species and are also widely used by humans (Beck et al., 2001; Barbier et al., 2010; Potter et al., 2015a). In recent years there has been growing recognition that the physical and biotic characteristics of estuaries with a small tidal range (*i*.*e*. microtidal; < 2 m), such as those in south-western Australia, are markedly different from those systems with a larger tidal range (Tweedley et al., 2016a; Rose et al., 2020). From a fish faunal perspective, the nearshore waters of microtidal systems are dominated by species capable of completing their life cycle within estuaries (*e*.*g*. atherinids and gobiids), with fish in offshore waters generally being marine estuarine-opportunists that use the estuary as a nursery area (Chuwen et al., 2009; Hoeksema et al., 2009). The benthic macroinvertebrate faunas also differ from macrotidal estuaries, predominantly comprising small deposit-feeding species and harbour insect larvae (Tweedley et al., 2016a,b; Warwick et al., 2020). Given their limited tidal water exchange and highly seasonal rainfall, south-western Australian estuaries are particularly vulnerable to degradation and climate change and thus require regular monitoring and management (Hallett et al., 2018; Warwick et al., 2018). While much focus has been placed on understanding the larger systems located near population centres, such as Perth, Mandurah and Bunbury (*e*.*g*. Veale et al., 2014; Potter et al., 2016; Valesini et al., 2018), less focus has been placed on smaller, more regional estuaries (Tweedley et al., 2017a; 2018).

Hill Inlet, *i*.*e*. the estuary of the Hill River, is a very small (0.06 km^2^) wave-dominated system located ∼200 km north of Perth. The estuary is ‘riverine’ in geomorphology and lies in Pleistocene sediments towards the northern extent of the Swan Coastal Plain (Hodgkin and Hesp, 1998). It is fed by Hill River, which is ∼80 km long, drains an area of 2,800 km^2^ and provides an average annual flow of 5,250 ML (Brearley, 2005). Hill River is relatively unique in this part of Western Australia, being the only such river that ‘meets’ the ocean in the 270 km between Dongara and Guilderton, with other rivers terminating in small fresh or saline lakes and swamps (Pen, 1999). Surveys in the 1960s found that water in this river was brackish, however, it is thought that prior to the clearing of 25% of the catchment for agricultural purposes, resulting in secondary salinization, that it was a freshwater system (Brearley, 2005). Like many of the microtidal estuaries in Western Australia, Hill Inlet does not maintain a permanent connection with the ocean, with longshore and onshore transport of sand forming a bar at the mouth of the estuary (Brearley, 2005; Hoeksema et al., 2018). This sand bar is reported to break each year (*i*.*e*. seasonally-open), the timing of which depends on when the first heavy and consistent rains occur. Typically, the bar will only remain open for a few days or weeks, with the duration influenced by the amount of fresh water within the estuary, which can scour away sand and influence sediment transport in coastal waters (Brearley, 2005).

In a classification of the ‘condition’ of 979 estuaries across Australia by the National Land and Water Resources Audit, Hill River was assigned the status of ‘modified’ (Commonwealth of Australia, 2002). This system, together with Toby Inlet and the Margaret River Estuary, were the only estuaries on the west coast of south-western Australia, not to be rated as ‘extensively modified’ (Tweedley et al., 2017b). However, the estuary and its catchments are been increasingly threatened by uncontrolled access by off-road vehicles, which, in other parts of the world, have been shown to have a range of impacts. These include accelerating dune erosion, damaging vegetation and crushing benthic invertebrates such as crabs, isopods and bivalves (Davenport and Davenport, 2006; Sheppard et al., 2009). The seriousness of the impacts of off-road vehicles led to South African authorities to ban such vehicles from large stretches of the KwaZulu-Natal coastline (Celliers et al., 2004). In addition to these direct impacts, off-road vehicles have an indirect influence by providing recreational fishers with access to sites that may have been previously inaccessible. Given the very small size of Hill Inlet, it is possible that if recreational fishing effort increased markedly as a result of additional visitors, fish populations could decline. For example, changes made to off-road vehicle access in South Africa led to a marked increase in recreational fishing in the Sundays Estuary and a decline in the stocks of recreationally-targeted fish species (Cowley et al., 2013).

In light of the above, and that fact that no baseline faunal data had been collected, the aim of this study was to provide the first quantitative description of the fish and benthic macroinvertebrate fauna of Hill Inlet. These data could then be used in management plans and form a benchmark against which future changes could be detected.

## Materials and methods

### Sampling regime

Fish fauna in the shallow, nearshore waters (< 1.5 m deep) of Hill Inlet were sampled during two days in June 2019 (austral winter), when the bar at the estuary mouth was closed to the ocean (Fig. 1). The initial aim was to sample along the longitudinal axis of the estuary at regular intervals in an upstream direction. However, due to high water levels only eight nearshore sites were able to be sampled, with all except one collected from the lower region of the estuary. Therefore, an additional 15 samples were collected from the most downstream areas that was shallower and able to be sampled effectively. Sampling in nearshore waters employed a seine net that was 21.5 m long and consisted of two 10 m long wings (6 m of 9 mm mesh and 4 m of 3 mm mesh) and a 1.5 m long bunt made of 3 mm mesh and swept an area of ∼116 m^2^. Upon capture, all fish were identified to species and returned to the water alive.

**Fig. 1.**
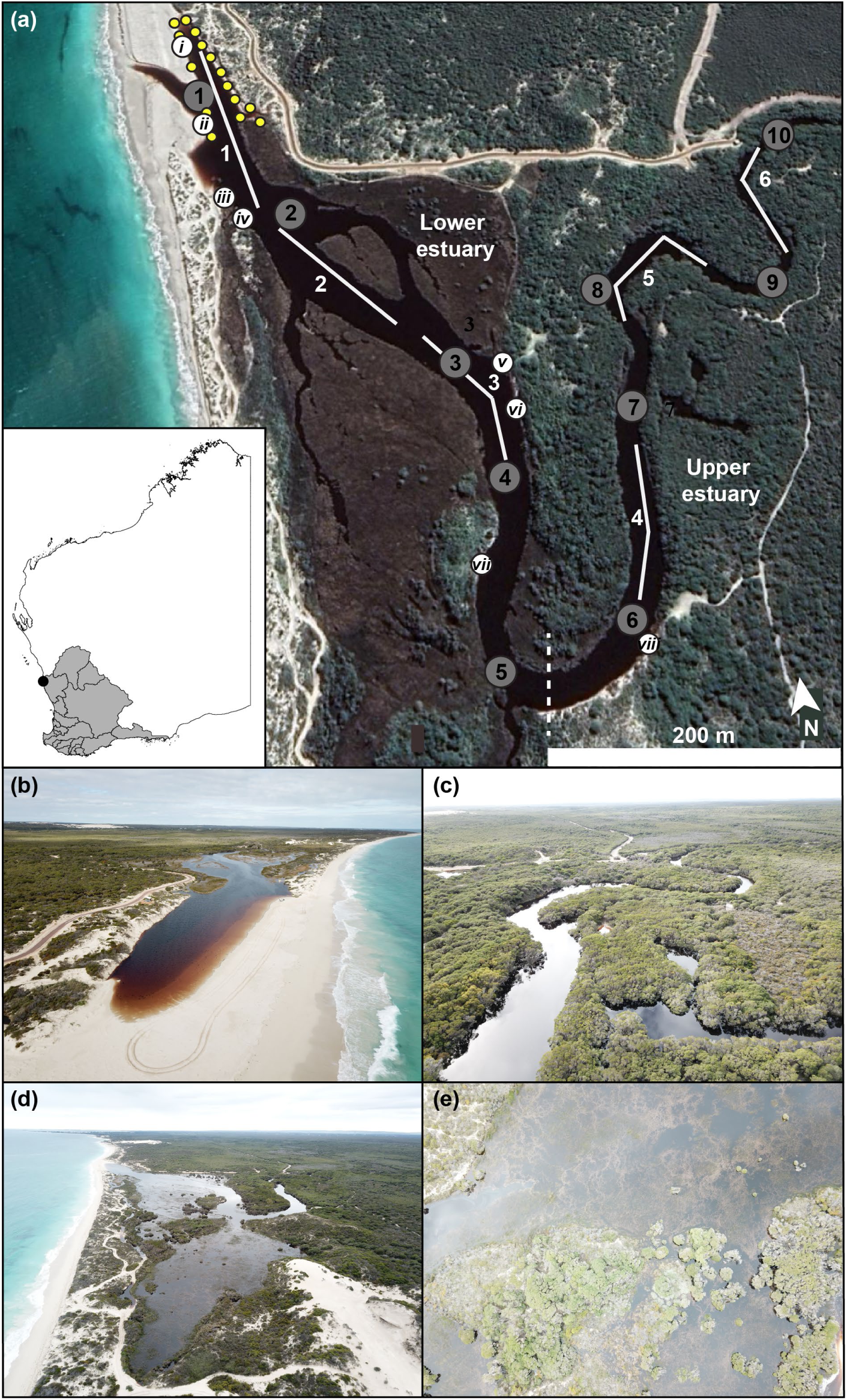
Satellite image of Hill Inlet (30°23’17.2”S; 115°03’09.3”E) taken in August 2018 showing (a) sites where seine nets (white circles) gill nets (white lines) and Ekman grab (grey circles) samples were collected. Note the water levels and lack of inundation of the surrounding salt marsh. Dashed white line denotes the division between the lower and upper estuary. Yellow circles show the sites in the downstream part of the lower region of Hill Inlet where an additional 15 seine net samples were collected. Location of Hill Inlet in Western Australia denoted by the black circle, grey shading shows the extent of the south-west drainage division. Photographs showing (b) lower and upper regions of Hill Inlet and (c and d) extent of the flooding of the surrounding salt marsh in June 2019. Satellite images provided by Google and CNES/Astrium.

Fish in the adjacent offshore waters (> 1.5 m deep) were sampled at three sites in each region, using a sunken composite multifilament gill net. This was the maximum number of nets that could be deployed without overlapping them in each region. The net, which has a total length of 160 m, comprised eight 20 m long panels, each with a height of 2 m and a different stretched mesh size, *i*.*e*. 35, 51, 63, 76, 89, 102, 115 or 127 mm. Gill nets were set for one hour, to minimise any fish deaths and allow the return of fish alive to the water (Grixti et al., 2010). Fish were identified to species, counted and measured to the nearest 1 cm before being released. Abundances were recorded as catch rate h^-1^.

Each fish species caught using either net was also assigned to a life-cycle guild within a category according to the way in which it uses estuaries (Potter et al., 2015b). Definitions of the two categories and two guilds relevant to the current study are as follows. **Marine category**, *i*.*e*. species that spawn at sea. *Marine estuarine-opportunist* (MEO) guild, *i*.*e*. species that spawn at sea and regularly enter estuaries in substantial numbers, particularly as juveniles, but can also use coastal marine waters as alternative nursery areas. **Estuarine category**, *i*.*e*. species with populations in which the individuals can complete their life cycles within the estuary. *Solely estuarine* (E) guild, *i*.*e*. species typically found only in estuaries. No species belonging to the **freshwater** or **diadromous** categories were recorded in this study.

Benthic macroinvertebrates were sampled at ten sites spread along the central longitudinal axis of the estuary using an Ekman grab operated from a boat, with five sites located in each of the upstream and downstream regions (Fig. 1). The grab collected sediment from an area of 225 cm^2^ and sampled to a depth of 15 cm. Each sediment sample from each site was wet-sieved through a 500 µm mesh in the field and preserved in a 5% formalin mixture buffered in estuary water. After a week the sediment was re-sieved and transferred to 70% ethanol. Using a dissecting microscope, any invertebrates found in a sample were removed from the sediment retained on the mesh and identified to the lowest possible taxonomic level. Water temperature (°C), salinity and dissolved oxygen concentration (mg L^-1^) were measured at the surface and bottom of the water column at each site where benthic macroinvertebrate samples were collected using a Yellow Springs Instrument 556 water quality meter (Yellow Springs Instrument, Ohio, USA).

### Statistical analyses

All univariate and multivariate statistical analyses were performed using SPSS v24 and the PRIMER v7 multivariate statistics software package (Clarke and Gorley, 2015), respectively. As the vast majority of the samples collected from the nearshore waters contained no fish (see Results), these data were not subjected to any statistical analyses.

The DIVERSE routine was used to calculate the number of species and total catch rate (fish h^-1^) in each gill net sample. The total catch rate was square-root transformed to meet the assumption of equal variance. Each variable was subjected to an Analysis of Variance (ANOVA) test to determine whether that variable differed among Region (2 levels; Lower and Upper estuary). In this and all other tests the null hypothesis of no significant difference among *a priori* groups was rejected if the test statistic (*P*) was < 0.05.

As only three gill nets were able to be deployed in each region and a minimum of four samples are required for multivariate analysis, the samples in each region were averaged to create a fourth replicate. The abundances of each species in each of the sample were square-root transformed to down-weight the contributions of species with high values in relation to those with lower values and used to construct a Bray-Curtis resemblance matrix. This matrix was subjected to a one-way Analysis of Similarities (ANOSIM) test (Clarke and Green, 1988). This same matrix was used to construct a non-metric Multidimensional Scaling plot to visualise the patterns of differences among samples. The square-root transformed abundances of each taxon in each sample were used to produce a shade plot. The resultant plot is a simple visualisation of the frequency matrix, where a white space for a taxon demonstrates that the taxon was never collected, while the intensity of grey-scale shading is linearly proportional to the abundance of that taxon (Clarke et al., 2014). The order of both the sites (*y* axis) and taxa (*x* axis) were determined by independent seriation. Thus, taxa exhibiting similar patterns of abundance across the regions were ordered together and *vice versa* for regions. These re-orderings are purely to aid visualisation and are irrelevant to multivariate analyses of those samples.

Data from the five replicate Ekman grab samples from each region were used to construct a data matrix of the number of each benthic invertebrate species (individuals 225 cm^-2^). This matrix was subjected to the DIVERSE, ANOVA, ANOSIM, nMDS and shade plot procedures described for the gill net samples, but without the need to create an average sample.

## Results

### Water quality

Surface water temperature declined essentially sequentially in an upstream direction from 15.3 to 13.6 °C. Values for this environmental variable at the bottom of the water column were fairly consistent at ∼17.2 °C at all except the three most downstream sites, where temperatures were between 15.3 and 16.4 °C (Fig. 2a). Salinity in the surface waters declined in an upstream direction from almost 12 near the bar at the mouth of the estuary to < 3 at the most upstream site, ∼1.5 km away (Fig. 2b). With the exception of site 1 where salinities were reduced (∼12), there was no conspicuous longitudinal change in salinity in the bottom waters (mean = 26.7). The salinity of 31 recorded at Site 10, does, however, highlight the penetration of the salt-wedge upstream.

**Fig. 2.**
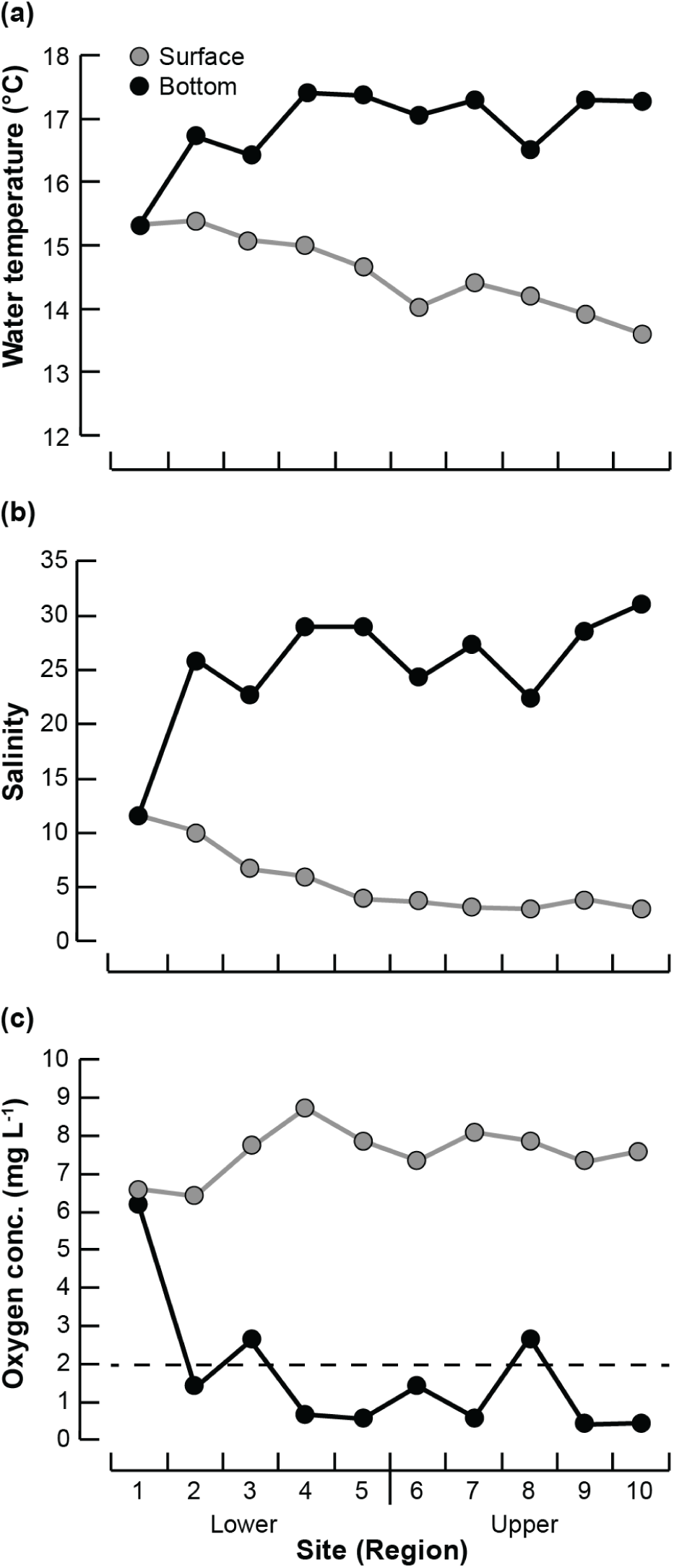
(a) Water temperature (°C), (b) salinity and (c) dissolved oxygen concentration (mg L^-1^) in the surface and bottom of the water column at each site in Hill Inlet. The dashed line in (c) denotes hypoxic conditions (< 2 mg L^-1^).

The typically oligohaline surface salinities were caused by the fact that the system is located in a Mediterranean region, where 80% of rainfall occurs between May and September (Hallett et al., 2018). Hill Inlet only received its first discharge from the Hill River for the calendar year on June 14 2019 (five days before sampling began) and 206 days after its last input of 0.003 ML on 19 November 2018 (Department of Water and Environmental Regulation, 2020). The 76 ML of flow that had occurred by the time sampling started was enough to add > 1 m of water over the 0.06 km^2^ area of the estuary, markedly lowering salinity in only the surface waters and causing pronounced inundation of the surrounding salt marsh. However, this flow was not sufficient to raise water levels enough to breach the sand bar at the mouth of the estuary (Fig. 1b-e), which would have increased salinity due to the tidal exchange of full-strength seawater.

Stratification index values (*i*.*e*. bottom salinity – surface salinity) were 0 at the well-mixed and shallow Site 1 and then increased with distance upstream to ∼16 at the next two sites and then always > 20 and reached as high as 28 at Site 10. This pronounced halocline at most sites likely caused the hypoxic conditions (< 2 mg L^-1^) present in the bottom waters of all except site 1 (Fig. 2c). A similar situation occurs in the Swan-Canning Estuary, a salt-wedge system located ∼230 km south, during times of intermediate flows, *i*.*e*. when there is not enough freshwater discharge to decrease the salinity of the entire water column, but enough to generate a halocline (*e*.*g*. Hamilton et al., 2001; Tweedley et al., 2016b). In contrast, normoxic and relatively high oxygen concentrations (*i*.*e*. 6.4 to 8.7 mg L^-1^) were recorded in the surface waters of each site (Fig. 2c).

### Nearshore fish fauna

The initial eight seine net samples yielded a single individual of the sparid *Acanthopagrus butcheri* (Site 4), with the only other faunal species found being 16 individuals of the caridean shrimp *Palaemonetes australis* recorded at Site 7. The fifteen additional samples conducted along the banks of the most downstream 200 m of the estuary returned two individuals of the gobiid *Pseudogobius olorum* and another *A. butcheri*. Therefore, in total, only four fish were collected from the 23 samples covering an area of ∼2,668 m^2^, *i*.*e*. one fish per ∼667 m^2^. This density is the lowest for any estuary in south-western Australia (Tweedley, 2011) and likely reflects the fact that water levels were at close to their peak before the bar would breach naturally, which occurred several days after our sampling. Under such conditions fish would be able to move into the surrounding salt marsh areas (Fig. 1d,e), which were too deep (> 2 m) to be sampled using the seine net. A similar situation was reported by Tweedley et al. (2014) in the Vasse-Wonnerup Estuary, where densities during spring and summer reached, on average, ∼800 fish 100 m^2^ when water levels are much lower and only 20-30 fish 100 m^2^ during winter, when the surrounding salt marsh habitat become inundated.

Aside from the low density of fish it was striking that no atherinids were collected. Individuals from this family are ubiquitous in south-western Australian estuaries and, together with gobies, can represent up to 99% of all fish recorded in nearshore waters (Hoeksema et al., 2009; Valesini et al., 2009; Hogan-West et al., 2019). Given their wide tolerances for salinity and presence in high abundances in all estuaries in the region (Potter et al., 1990; Tweedley et al., 2019b), it is hypothesised that their absence in the current study was related to the elevated water levels and inundation of surrounding salt marsh. Furthermore, as the deeper waters were largely hypoxic, a vertical movement into the water column would be advantageous (à la Poh et al., 2019). Moreover, the structural complexity provided by the surrounding flooded marsh would limit predation from larger piscivorous fish (Humphries et al., 1992), such as the pomatomid *Pomatomus saltatrix* (see later). Both of the fish species recorded are solely estuarine species, that are able to complete their lifecycle within the estuary (Gill et al., 1996; Beatty et al., 2018), which fits the trend for this guild to dominate in systems that do not maintain a permanently-open connection to the ocean (Hoeksema et al., 2009; Tweedley et al., 2016a).

### Offshore fish fauna

Gill netting in the offshore waters of Hill Inlet yielded 344 fish, representing four species and three families (Table 1). The Mugilidae was the only family to comprise two species. In contrast to the nearshore waters, only one of the species is able to spawn in estuaries (*i*.*e. A. butcheri*), with the remaining three species all being marine estuarine-opportunists. A moderate size range (> 170 mm) of individuals from each species except *P. saltatrix* were recorded, suggesting there were multiple ages classes present in the estuary (Chubb et al., 1981; Cottingham et al., 2015). *Acanthopagrus butcheri* and two species of mullet, comprised the vast majority of individuals (99%), which is fairly common in small estuaries on the west coast of Western Australia (Tweedley et al., 2018; Cottingham et al., 2019).

**Table 1.** Mean catch rate (h^-1^; *X*), standard error (SE), percentage contributions to the total catch (%), ranking by contribution (R), mean total length (L, mm) and length range (Lr) of each fish species caught in the offshore waters of Hill Inlet in June 2019. Those species that contributed ≥ 10% of the total catch are shaded in grey. The estuarine usage functional group (EUFG) to which each species belong is provided (E, solely estuarine; MEO, marine estuarine-opportunist). Species ranked by total number of fish caught (n).

| Scientific name | Family | EUFG | n | $X$ | SE | % | R | L | Lr |
| --- | --- | --- | --- | --- | --- | --- | --- | --- | --- |
| *Mugil cephalus* | Mugilidae | MEO | 167 | 27.83 | 6.51 | 48.55 | 1 | 292 | 120-400 |
| *Acanthopagrus butcheri* | Sparidae | E | 140 | 23.33 | 5.25 | 40.70 | 2 | 196 | 130-300 |
| *Aldrichetta forsteri* | Mugilidae | MEO | 35 | 5.83 | 2.25 | 10.17 | 3 | 279 | 155-500 |
| *Pomatomus saltatrix* | Pomatomidae | MEO | 2 | 0.33 | 0.00 | 0.58 | 4 | 360 | 350-370 |

The lack of common local species, such as the terapontids *Amniataba caudavittata* and *Pelates octolineatus*, the sparid *Rhabdosargus sarba* and the sillaginids *Sillago schomburgkii* and *Sillaginodes punctata*, is likely to reflect the limited number of samples collected and the fact that the bar at the mouth of the estuary was closed and had been for some time due to the lack of freshwater discharge and along-shore transport of sediment. In Toby Inlet, a small estuary, located ∼400 km south of Hill Inlet, the number of species recorded from 16 gill net samples increased from four when the bar was closed to 12 when the bar was open, with the ‘additional’ species all being regarded as marine, *i*.*e*. marine estuarine-opportunists and marine stragglers (Tweedley et al., 2018).

Despite the invasive Goldfish (*Carassius auratus*) and Mozambique Tilapia (*Oreochromis mossambicus*) having been recorded in the Moore and Chapman rivers, respectively (Morgan et al., 2004), which lie to the south and north of Hill Inlet neither were recorded in the current study. As both species are able to survive in saline waters (Whitfield et al., 2006; Beatty et al., 2017), there is value in being alert to the presence of these species in any future surveys. An individual of the chelid turtle *Chelodina colliei* was caught at each of sites 3 and 5 and returned live to the water.

The mean number of fish species in offshore waters differed significantly among regions (F_1, 1_ = 16.00, *p* = 0.016) being slightly greater in the lower (3.3) than upper (2.0) estuary (Fig. 3a). However, there was no significant difference in mean catch rate between regions (F_1, 1_ = 1.84, *p* = 0.246; Fig. 3b). This was due to the large variability among the replicate samples from the upper estuary with catches of 68, 54 and only 12 fish h^-1^. ANOSIM detected a significant and moderate difference in fish faunal composition (*P* = 0.029; Global *R* = 0.543), with the points for each region in offshore waters forming discrete groups on the nMDS plot (Fig. 4).

**Fig. 3.**
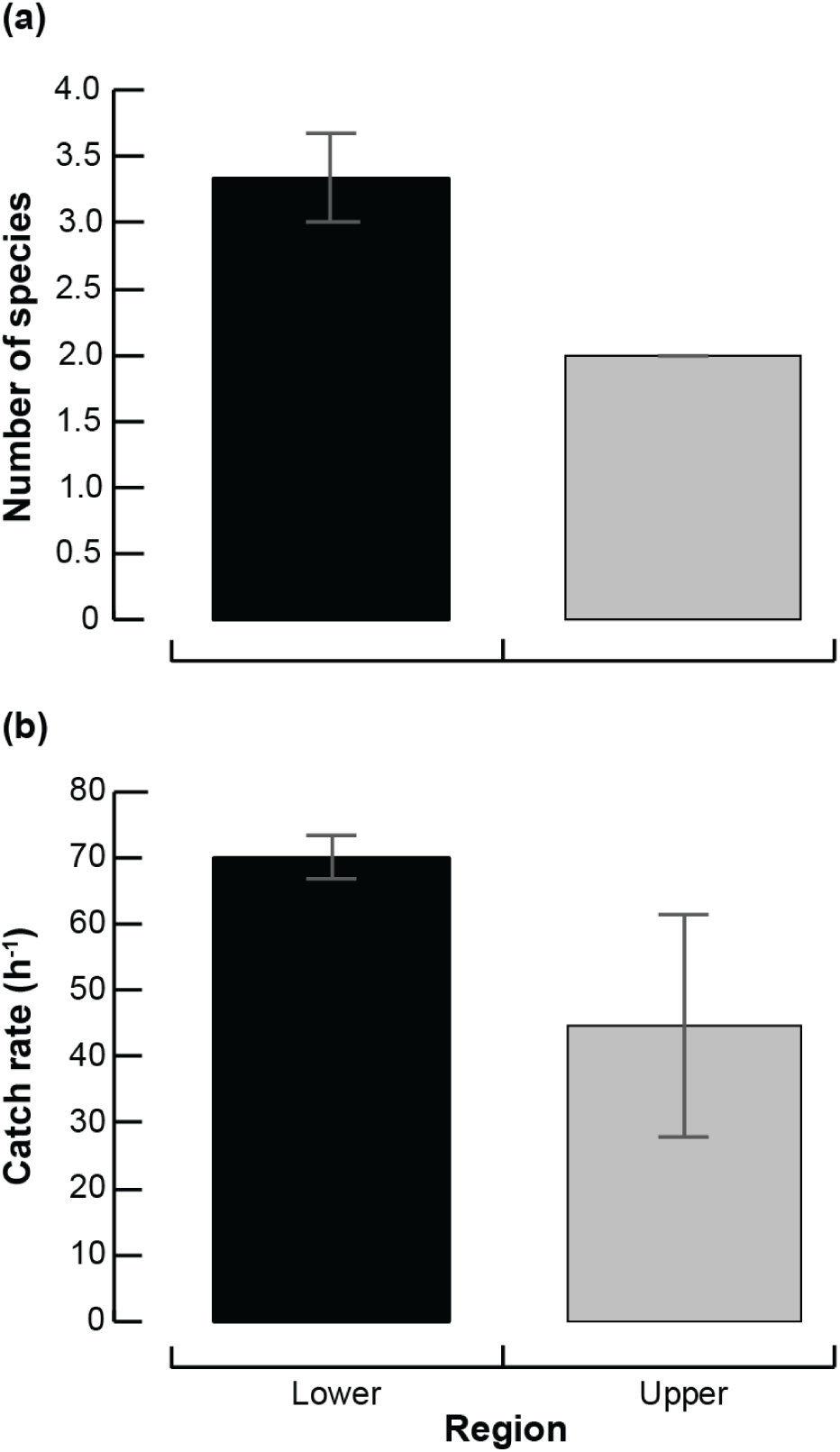
Mean (a) number of fish species and (b) catch rate (fish h^-1^) caught using gill nets in each region of Hill Inlet. Error bars represent ± 1 standard error.

**Fig 4.**
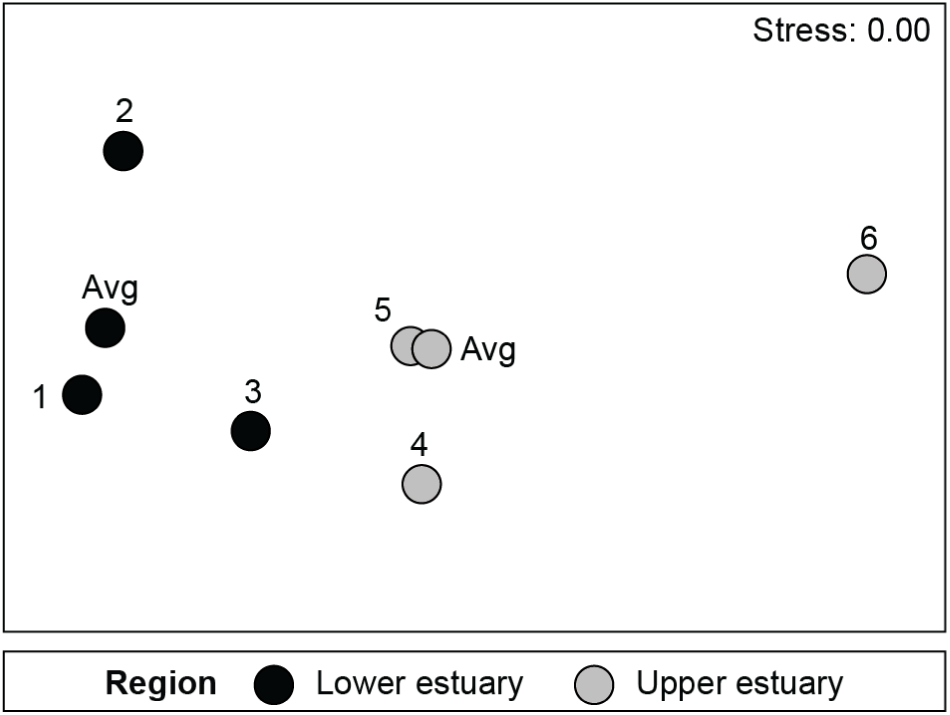
Two-dimensional nMDS ordination plot constructed from a Bray-Curtis resemblance matrix of the square-root transformed catch rate (h^-1^) of each fish species in each gill net sample from the offshore waters of the Hill Inlet.

Although *M. cephalus* and *A. butcheri* dominated the catches at all sites, there was a longitudinal shift in faunal composition, with *P. saltatrix* only being found in the most downstream site, and the mugilid *Aldrichetta forsteri* only at sites in the lower estuary (Fig. 5). This accounts for the number of species recorded per sample declining from four at the bar, to three in the remaining sites in the lower estuary and two at each site in the upper estuary. This change in fish species along the estuary reflects the fact that *A. butcheri* is a solely estuarine species, which resides in upper reaches and is highly euryhaline and that, although *M. cephalus* is a marine species, when present in estuaries, individuals are most typically abundant in upstream regions (Loneragan et al., 1989; Chuwen et al., 2009). In contrast, *P. saltatrix*, is typically found in more saline conditions in the lower reaches of the estuary. In this study, while more saline conditions were found in the deeper waters of the upper estuary, these individuals may be responding to cues of marine water seeping through the bar and preparing to leave the estuary immediately upon the bar breaking, which occurred several days post sampling.

**Fig. 5.**
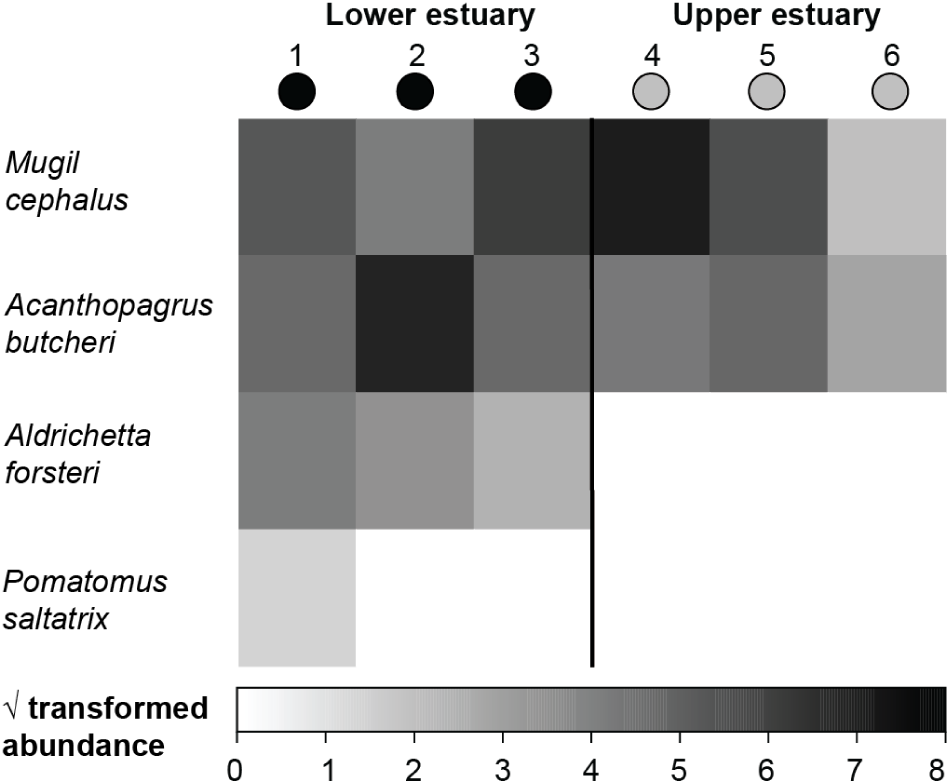
Shade plot of the square-root transformed catch rate (h^-1^) for each fish species recorded in the offshore waters of each site in Hill Inlet. White space indicates that a species was absent, and the shades of grey are proportional to the transformed abundances.

### Benthic macroinvertebrates

A total of 631 benthic macroinvertebrates were recorded from the 10 Ekman grab samples collected from the sediment in the offshore waters of Hill Inlet (Table 2). These invertebrates comprised 10 species from four phyla. While the molluscs were the most speciose group constituting three species, members of this group contributed 30 % to the total number of individuals, which is less than the two species of polychaete that represented 47 % of the total number of benthic invertebrates (Table 2). The top five species all contributed > 5 % to the total number of individuals, together comprising 97% of the total numbers. Among these the polychaete *Capitella capitata*, bivalve mollusc *Fluviolanatus subtorta* and crustacean Ostracod sp. were the most abundant taxa (Table 2). The dominance of polychaetes, molluscs and crustaceans in Hill Inlet is typical for estuaries both in south-western Australia and more generally around the world (Tweedley et al., 2012; 2015). Recent work in Toby Inlet and the Vasse-Wonnerup, both of which are similar to Hill Inlet in terms of their relatively small size, presence of fringing salt marsh, oligohaline waters and limited connectivity to the ocean has shown the speciose nature of their insect fauna, with some of those taxa being relatively abundant (Tweedley et al., 2018; 2019a). Given the abundance of insects in these similar systems, it was surprising not to see more of these taxa here, particularly during times of the year when salinities are lower. This could reflect the limited number of samples collected and the snapshot nature of the study (*i*.*e*. all samples obtained over two days).

**Table 2.**
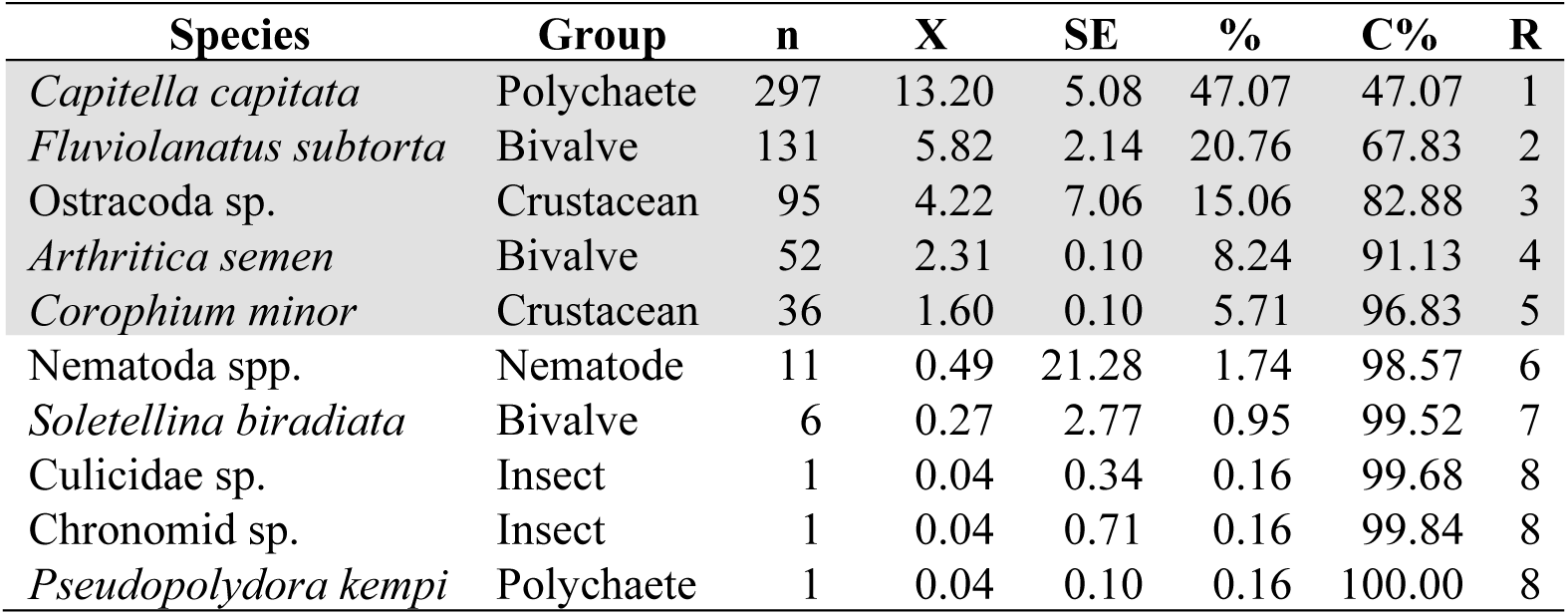
Mean density (100 cm^-2^; X), standard error (SE), percentage contributions to the total catch (%C) and ranking based on abundance (R) of each benthic macroinvertebrate species caught in the offshore waters of Hill Inlet in June 2019. Those species that contributed ≥ 5% of the total catch are shaded in grey. Species ranked by total number of individuals caught (n). The broad taxonomic group to which each species belongs is also provided.

The mean number of benthic macroinvertebrate species differed among regions (F_1, 8_ = 11.25, *p* = 0.029), being greater in the lower (5) than upper (2) estuary (Fig. 6a). Despite the mean number of individuals being four times greater in the lower estuary (Fig. 6b), it did not differ significantly among regions (F_1, 8_ = 1.84, *p* = 0.243). This was due to the large variability among the replicate samples from the lower estuary (range = 9 to 340; Fig 6b). Note that no benthic invertebrates were recorded in the sample from the upper most site. There was, however, a significant and moderate difference in faunal composition between the two regions (ANOSIM *P* = 0.008; Global *R* = 0.508). This is clearly shown on the nMDS plot where there is a complete separation of points representing the samples from the lower and upper estuary (Fig. 7). In terms of the species responsible for the differences, the microbivalve *Arthritica semen*, polychaete worm *C. capitata*, the sunset shell *Soletellina biradiata* and nematode spp. typified the lower estuary and were not recorded in the upper sites (Fig. 8). The only species found in more than one of the sites in the upper estuary were the bivalve *F. subtorta* and the amphipod *Corophium minor*, although these species were also recorded at multiple sites in the lower estuary. The presence of these species throughout the entire estuary reflects their ability to tolerate a wide range of salinities (Morton, 1982; Tweedley, 2011). While it is hard to draw conclusions from the small range of samples collected here at one point in time, the relatively low diversity (cf. this study; Tweedley et al., 2019a) could be due to the hypoxia present throughout much of the deeper waters (Fig. 2c). Hypoxia influences the diversity and composition of benthic fauna, reducing the number of species, particularly crustaceans (Diaz and Rosenberg, 1995; Gray et al., 2002; Tweedley et al., 2016b).

**Fig. 6.**
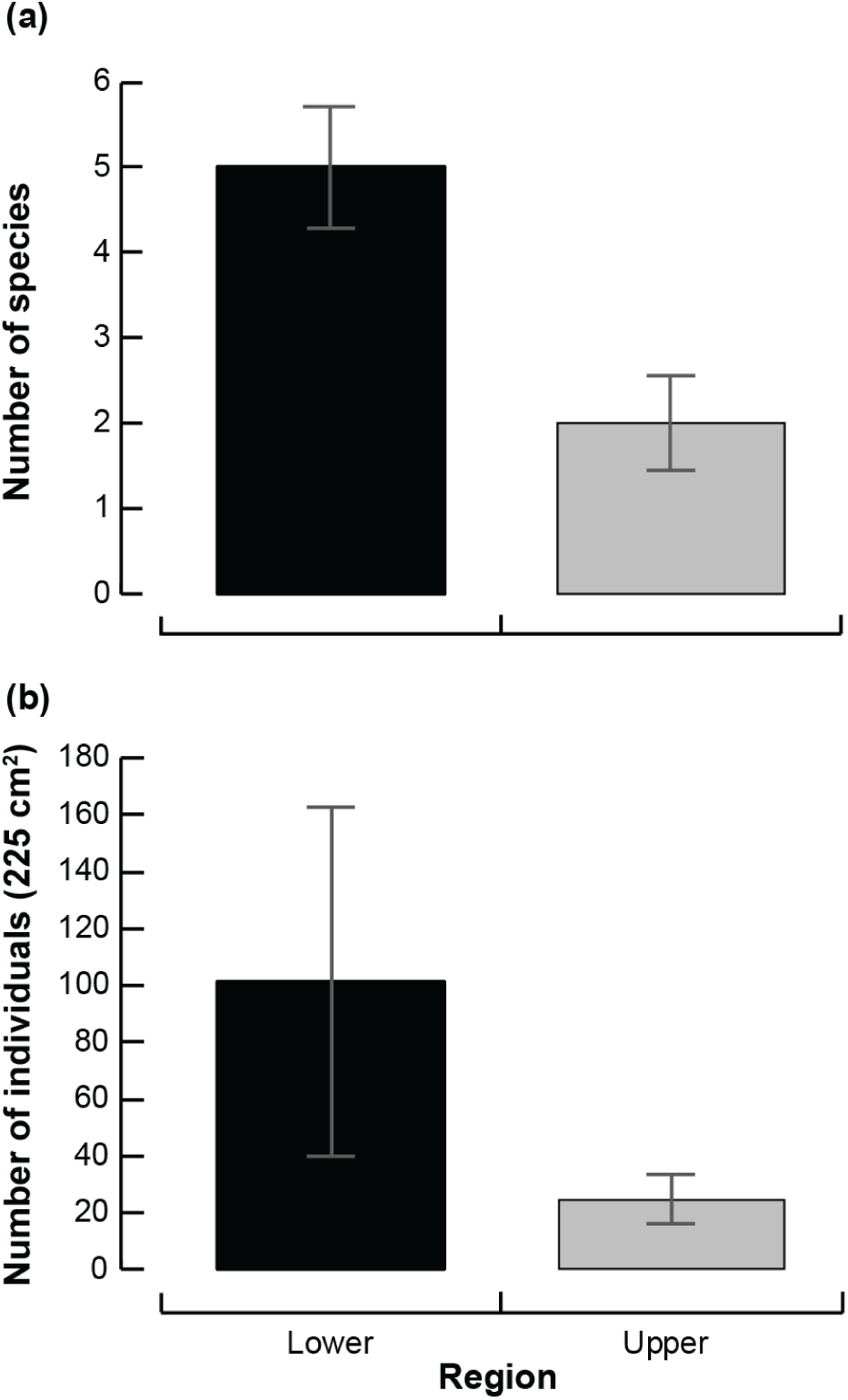
Mean (a) number of benthic macroinvertebrate species and (b) individuals (225 cm^2^) in each region of Hill Inlet. Error bars represent ± 1 standard error.

**Fig 7.**
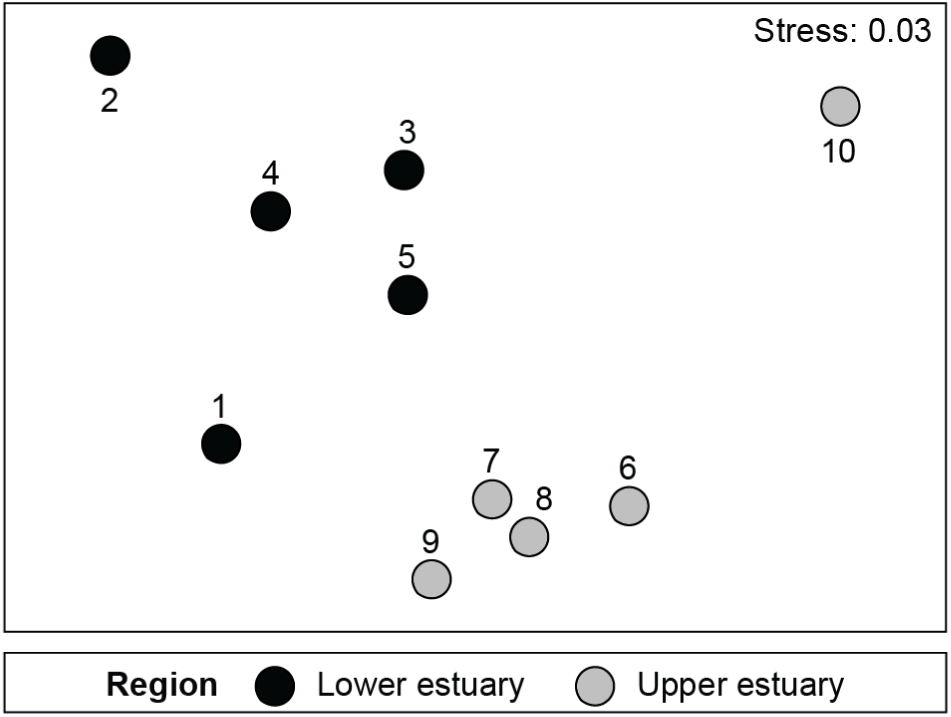
Two-dimensional nMDS ordination plot constructed from a Bray-Curtis resemblance matrix of the square-root transformed abundance of each benthic macroinvertebrate species in each Ekman grab sample from the two regions of Hill Inlet.

**Fig. 8.**
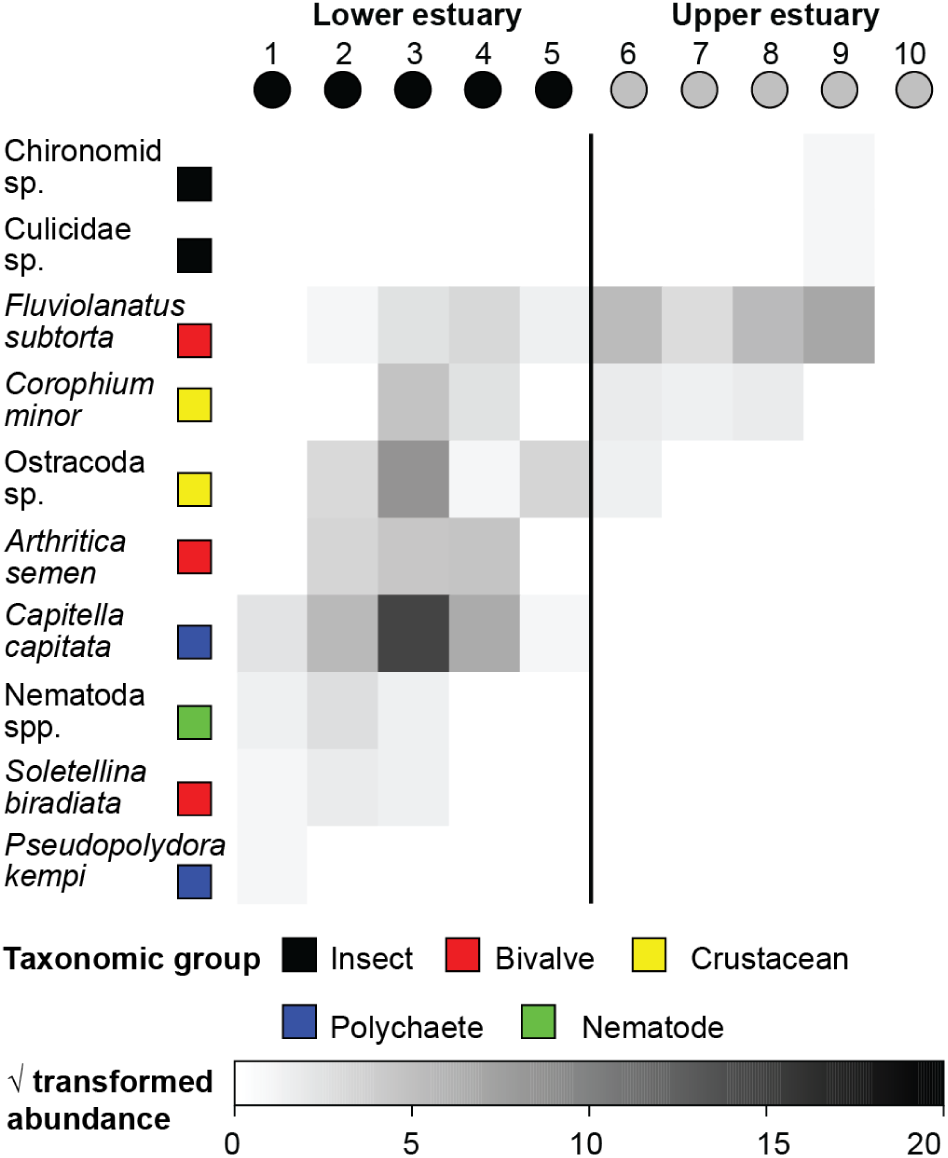
Shade plot of the square-root transformed abundance (225 cm^-2^) for each benthic macroinvertebrate species in each Ekman grab sample from the two regions of Hill Inlet. White space indicates that a species was absent, and the shades of grey are proportional to the transformed abundances. The broad taxonomic group to which each species belongs is also provided.

## Conclusions

This study obtained the first quantitative data on the fish and invertebrate fauna of Hill Inlet a very small, seasonally-open estuary located on the norther most part of south-western Australia. Relatively few fish species were recorded, which is likely to reflect the high water levels allowing fish to immigrate into surrounding salt marsh, and the bar at the mouth of the estuary being closed, thus preventing recruitment of marine species. The species that were recorded are common in south-western Australian estuaries, with two able to complete their entire life cycle within Hill Inlet. The benthic fauna of Hill Inlet was fairly depauperate and likely impacted by the presence of hypoxia throughout much of the bottom waters. As the data in this study were collected over a small temporal window sampling should be repeated at a time of the year when water levels are lower, either during summer or again in winter, but only after the bar has broken to provide a more holistic assessment of the fauna of Hill Inlet. In the meantime, given the lack of scientific information available on Hill Inlet, there would be value in engaging with the community, such local ecological knowledge has been shown to be useful in informing decision making and management (Schlacher et al., 2010; Obregón et al., 2020).

## Acknowledgements

The authors gratefully acknowledge the support of Dr Nic Dunlop and Alison Goundrey from the Conservation Council of Western Australia and people from the Tending the Tracks Alliance and the Subaru 4WD Club of Western Australia for the great assistance with the fieldwork. Kurt Krispyn is thanked for helping to process the benthic macroinvertebrate samples. Funding was provided by the Northern Agricultural Catchments Council, with support from the Western Australian Government’s Natural Resource Management Program. All work was carried out in accordance with Murdoch University Animal Ethics Permit R2987/17 and Department of Primary Industries and Regional Development Exemption #2987.

